# Whole genome assembly and annotation of the clover root weevil (*Sitona obsoletus*) using a combination of Illumina, 10X Genomics and MinION sequencing

**DOI:** 10.1101/2023.10.03.560759

**Authors:** Mandira Katuwal, Craig B. Phillips, Neil J. Gemmell, Eddy Dowle

## Abstract

Weevils are a highly diversified taxon, comprising about 70,000 described species that include many agricultural pests, biological control agents, and nutrient recyclers. Despite their importance and vast diversity, the number of sequenced genomes for the weevil family is still low (n=15). Here, we present a high-quality and contiguous genome assembly of *Sitona obsoletus* (Coleoptera: Curculionidae: Entiminae), a widespread invasive forage pest that infests clover species (*Trifolium* spp.) worldwide. We sequenced, assembled, and annotated the *S. obsoletus* genome using a hybrid approach that employed Nanopore long reads, 10X Chromium linked reads, Illumina short reads for assembly, and mRNA short read sequencing of various developmental stages for annotation. Our final annotated genome has a size of 1.2 Gb, with an N50 length of 313.85 kb. Benchmarking analyses against conserved single-copy Orthologs (BUSCO) found that over 94% of the genes were complete from each of the three BUSCO databases (Eukaryota, Insecta, and Arthropoda). A total of 9,777 protein-coding genes were annotated using the MAKER2 pipeline, of which 65% were functionally annotated. The annotated repeat elements make up 84.26% of the genome. The high-quality annotated genome of the weevil will facilitate a wide range of genetic, genomic, and phylogenetic studies on invasive weevils, as well as other weevil species in the subfamily Entiminae.

## Introduction

Weevils (Coleoptera: Curculionidae) constitute a mega-diverse group of beetles with about 70,000 described species (Alonso-Zarazaga, Barrios et al. 2017). They are among the largest family of animals described and inhabit nearly all terrestrial habitats. Being highly diversified taxon, they serve as agricultural pests, nutrient recyclers, pollinators, and often natural enemies (McKenna, Sequeira et al. 2009). Given their ecological and economic importance, genome sequencing of weevil pests are of great interest (Van Dam, Cabras et al. 2021). However, only 15 species from the family Curculionidae have publicly available genomes (Mei, Jing et al. 2022). The genus *Sitona* (Germar, 1817; Coleoptera: Curculionidae) comprises approximately 100 species, many of which are significant agricultural pests (Petrukha 1970, Alonso-Zarazaga 2007, Park, McNeill et al. 2013, Sanaei, Seiedy and de Castro 2015). Among them, clover root weevil *Sitona obsoletus* (Gmelin,1790; Coleoptera: Curculionidae: Entiminae) is a destructive clover (*Trifolium* spp.) pest.

*Sitona obsoletus*, an obligate clover feeder, is of European origin (Basse, Phillips et al. 2015), but has spread to numerous countries across Asia (Coşkuncu and Gencer 2010, Seebens, Blackburn et al. 2017), North America (Bright 1994) and Oceania (Barker, Addison et al. 1996). The larvae are the most destructive stage as they feed on the plant roots and root nodules, leading to reduced nitrogen fixation followed by decrease in plant yield and survival (Gerard, Hackell and Bell 2007, Basse, Phillips et al. 2015). The adult weevil is highly dispersible due to its strong flying and hitchhiking ability. An individual female can lay over 1,000 eggs. Although typically univoltine (Gerard, Goldson et al. 2010), some invasive *S. obsoletus* populations, such as those in New Zealand are bivoltine (Gerard, Crush et al. 2020). Due to its invasive nature and high dispersal ability, *S. obsoletus* can establish itself quickly in a new region, causing significant economic losses (Gerard, Crush et al. 2020). For example, since its introduction in New Zealand in 1996 (Barratt, Barker and Addison 1996), it has remained one of the significant clover pests, costing 235 million NZD each year to pastoral agriculture (Ferguson, Barratt et al. 2019).

Heavy reliance on biological control measures without integrating other management approaches may lead towards the evolution of host resistance against parasites (Mills 2017, Tomasetto, Tylianakis et al. 2017, Tomasetto, Cianciullo et al. 2018). There has been increasing interest in developing more effective genetic pest control strategies globally (Dearden, Gemmell et al. 2018). A comprehensive understanding of the genetic make-up of the target species is essential to develop effective genetic pest control strategies. However, in the case of *S. obsoletus*, the lack of annotated genome assembly has hindered progress in the field. To address this, we assembled the whole genome of *S. obsoletus* and utilised transcriptomic data to annotate the genome obtained from the weevil’s various development stages and adult tissues. The resulting annotated genome will provide the basis for conducting comparative genomic studies and will aid in developing novel genetic control strategies within the sub-family Entiminae and other curculionids.

## Materials and Methods

### Insect sampling

*Sitona obsoletus* adults were collected from commercially farmed fields of white clover (*Trifolium repens* L.) and ryegrass (*Lolium perenne* L.) using a modified vacuum suction device at three locations in the South Island of New Zealand: Invermay (-45.85626, 170.38772), Rakaia Island (-43.845, 172.177), and Lincoln (-43.64230, 172.47090). The weevil eggs used for transcriptome sequencing were obtained from adult weevils collected from Lincoln. Adult weevils were cleaned and snap-frozen in liquid nitrogen and stored at -80°C until further processing. To determine parasitism status, each weevil was dissected in 1xPBS buffer under a dissection microscope following the protocol of Goldson and Emberson (1981) and tissues obtained from a single unparasitised weevil were used for subsequent nucleotide extraction.

### Illumina short-read sequencing and library preparation

Genomic DNA (gDNA) was extracted from a whole adult female *S. obsoletus* using a modified protocol based on Gemmell and Akiyama (1996), substituting LiCl with NaCl as described by (Al-Jiab, Gillum et al. 2019). DNA quality and quantity were assessed using a Nanodrop and a Qubit 2.0 Flurometer (Life Technologies, USA). A PCR-free library with an insert size of approximately 450 bp was constructed using an Illumina TruSeq DNA PCR-Free kit following the manufacturer’s protocols. The library thus constructed was sequenced on a single rapid run lane of the Illumina HiSeq2500 to generate 250 base pairs (bp) paired-end reads. Sequencing was conducted at the Otago Genomics Facility (OGF), University of Otago, Dunedin, New Zealand, following standard Illumina protocols.

### 10X Genomics linked-read sequencing and library preparation

DNA was extracted from a single male individual following the 10X genomics protocol for gDNA extraction for a single insect (https://support.10xgenomics.com/permalink/7HBJeZucc80CwkMAmA4oQ2). Briefly, dissected tissues were hand homogenized using a sterile homogenizing pestle in a 2ml Eppendorf tube then incubated at 37°C in 600 μl of lysis buffer (10 mM Tris-HCl, 400 mM NaCl, and 100mM EDTA, pH 8.0) with 100 μl of Proteinase K (20 mg/ml) for overnight digestion. gDNA was obtained after salting out by adding 240 μl 5 M NaCl and cleaning using 70% ethanol. Extracted DNA was quantified, and quality checked using the Qubit 2.0 Flurometer (Life Technologies, USA) and Nanodrop respectively. DNA with a size of over 40 kb was selected using a Blue Pippin (Sage Science, USA) and used for library preparation and sequencing. A standard 10X linked read library preparation was performed on the size selected DNA at the Genetic Analysis Services (GAS), University of Otago, Dunedin, New Zealand. The pooled library was sequenced on the Illumina Novaseq platform to generate 2x151 bp paired-end reads at Garvan Institute, Australia. However, the data yield was insufficient to achieve the suggested 56x coverage for the Supernova assembler. Therefore, the same library was re-sequenced on a single lane of the rapid flow cell of HiSeq 2500 at OGF to increase the raw yield.

### Oxford Nanopore long-read sequencing and library preparation

Five sequencing libraries were prepared using the ligation sequencing kit (SQK-LSK109) from Oxford Nanopore Technologies Ltd, Oxford, UK. The first two libraries were prepared from the gDNA isolated from the head and leg tissues of an individual weevil of unknown gender using the DNeasy Blood and Tissue Kit (Qiagen, Germany) following the manufacturer’s instructions. The step was followed by an AMPure XP bead clean up in the ratio 1.8:1 of beads and DNA respectively. The input DNA for the both libraries were ∼0.5 μg and the libraries were individually run on a single flow cell (FLOW-MIN106D) on the MinION sequencer with R9.5 chemistry operated with MinKNOW (version 20.06.4). The first library was sequenced for 24 hours, and then the flow cell was flushed using a Flow cell wash kit (Oxford Nanopore Technologies, Oxford, UK), followed by the loading, and running of the second library until pore exhaustion. The remaining three libraries were prepared using gDNA obtained using the tissues from whole-body of three adult individuals (two males and one female), with a range of 1.4-3.4 μg of gDNA following the 10X protocol described earlier. The extracted DNA was subjected to an AMPure XP bead clean-up step (ratio 1.8:1) and five times needle sheared (26G needle) to prevent clogging of sequencing pores. Each of the three libraries were sequenced on a single flow cell (FLOW-MIN106D) on the MinION sequencer with R9.5 chemistry for 72 hours on MinKNOW (version 20.06.4). Real-time base calling was turned off during sequencing and was performed later using the high-performance computing platform for all the five MinION runs, as described in the assembly strategy.

### RNA extraction and sequencing

Weevil eggs were collected and placed in an artificial diet on 24 well-plates and reared under laboratory settings at room temperature (18-20°C). Total RNA was extracted from whole individuals of different development stages (larvae and pupae of unknown sexes) and dissected tissues (head, abdomen, gonads) from adult unparasitised males and females using a Direct-zol RNA Microprep kit (Zymo Research, USA), including DNAse treatment following the manufacturer’s instructions. RNA isolation was performed separately for each tissue and developmental stage sample. The extracted RNA was then checked for quantity, quality and intactness using a Qubit 2.0 Flurometer, Nanodrop and 0.5% agarose gel respectively.

RNA integrity was further evaluated using a Fragment Analyzer (Advanced Analytical Technologies, Inc. USA) at OGF. Insects can exhibit variable RNA quality numbers (RQN) due to collapsing of 28S peak (Winnebeck, Millar and Warman 2010). The RQN values obtained from the 23 samples were ranged from 3.1-9, where values above seven are considered high quality, and 19 samples had RQN values greater than seven. For samples with an RQN less than seven, the trace was used to evaluate the RNA quality. All 23 samples that passed the quality controls were subjected to prepping using an Illumina Truseq stranded mRNA library kit before being pooled at equimolar concentration. The prepared pool was sequenced on a single lane of HiSeq 2500 V2 Rapid Sequencing (2*150 bp paired-end) at OGF.

### Genome size estimation

*Sitona obsoletus* genome size was estimated using the k-mer counting method and flow cytometry (Pflug, Holmes et al. 2020). For k-mer methods, Jellyfish v.2.3 (Marçais and Kingsford 2011) was used on Illumina shotgun sequencing data after Trimmomatic v 0.39 quality control step using the parameters (ILLUMINACLIP: TrueSeq3-PE.fa:2:30:10 SLIDINGWINDOW:4:20 LEADING:10 TRAILING:10 MINLEN:35) to generate a 21-mers k-mer frequency distribution. The resulting output was uploaded to GenomeScope v.2.0 (Vurture, Sedlazeck et al. 2017) to generate histograms for estimating the parameters such as genome size, repeat content and heterozygosity, with k-mer size set at 21, read length at 251 and max k-mer coverage at 1000.

In addition, the genome size was also estimated using flow cytometry using two biological replicates at Flowjoanna (Palmerston North, NZ), similar to the methodology of (Katuwal, Bhattarai et al. 2022). Briefly, an individual weevil head was detached and homogenized using a pestle in 500 μl of buffer solution containing 0.1% weight/volume trisodium citrate dihydrate, 0.1% volume/volume IGEPAL, 0.052% weight/volume spermine tetrahydrochloride, and 0.006% sigma 7-9 (all from Sigma-Aldrich, USA). Rooster red blood cells (RRBC) from a local rooster stored in citrate buffer were used as the reference sample. Test samples were filtered using a 35 μl filter cap before adding 100 μl of 0.21 mg/ml trypsin for further dissociation, and 75 μl of 2.5 mg/ml trypsin inhibitor (both from Sigma-Aldrich) for 10 minutes at 37 °C. Nuclei staining was performed using 100μl of pre-stain containing propodium iodide 416 mg/ml and RNAse 500 mg/ml. One sample was prepared with RRBC and the other without it. Both samples were measured using a FACSCalibur (BD Biosciences, USA) set with a 488 nm laser for producing fluorescence. This inclusion of RRBC facilitated to distinguish RRBC nuclei from the weevil’s nuclei. Thus, obtained data were further analysed using Flowjo (BD Biosciences, USA). The final size calculation was done using the pg/nuclei of the weevil samples.

### Bioinformatic pipeline

All bioinformatics analyses were completed using the New Zealand eScience Infrastructure (NeSI). The pipeline used in the study is described below (Figure 1).

**Figure 1.**
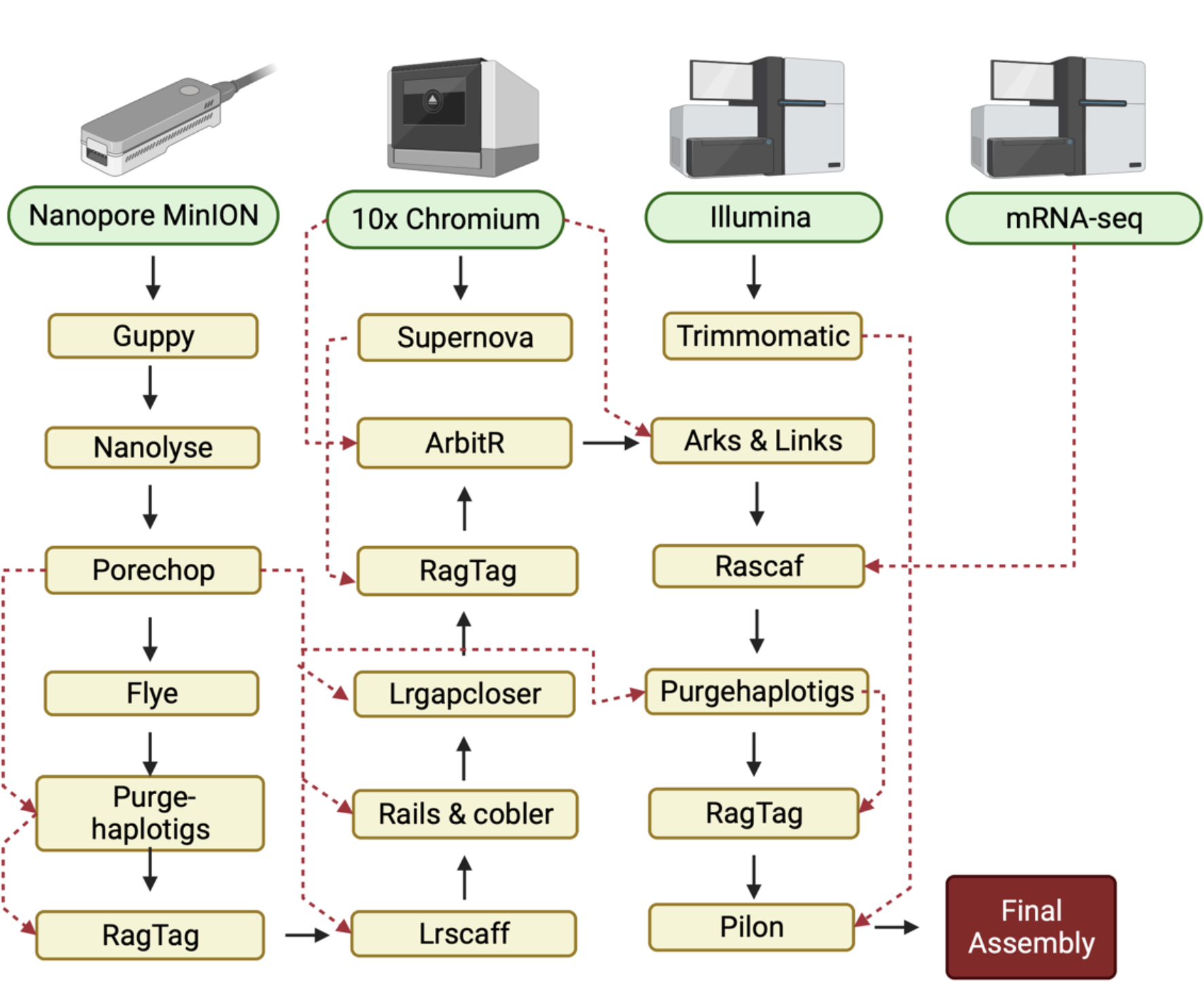
Schematic representation of the assembly pipeline for the *Sitona obsoletus* genome. The black arrow represents the workflow, and the red dotted line represents the additional input data in the pipeline (Created with Biorender.com).

### Assembly strategy

The quality of the raw data from the shotgun Illumina paired-end sequencing was evaluated using FastQC v 0.11.9 (Andrews 2017). The reads were then processed to remove the adapters and low-quality bases using Trimmomatic v 0.39 with options: ILLUMINACLIP: TrueSeq3-PE.fa:2:30:10 SLIDINGWINDOW:4:20 LEADING:10 TRAILING:10 MINLEN:35. The resulting short reads were subjected to *de novo* assembly using ABySS v.2.2.5 (Simpson, Wong et al. 2009).

The 10X Chromium linked reads were assembled using Supernova assembler v2.1.1 (Weisenfeld, Kumar et al. 2017). To obtain a haploid assembly, the final fasta sequence from the Supernova assembly was exported with the ‘pseudohap’ style. The assembly completeness was measured using BUSCO v.5.2.2 (Simão, Waterhouse et al. 2015), which compares single-copy orthologous genes. In addition, Quast v 5.0.2 (Gurevich, Saveliev et al. 2013) was used to generate other assembly metrics, such as N50, and number of contigs/scaffolds for evaluating overall contiguity of the raw and subsequent processed assemblies.

Raw fast5 Nanopore sequencing reads were basecalled using Guppy v.5.0.7 (Vereecke, Bokma et al. 2020) and pycoQC v.2.5.2 (Leger and Leonardi 2019) was used to access data quality. We used NanoLyse v.0.5.1. (De Coster, D’Hert et al. 2018) to remove library control lambda phase genome DNA (DNA CS). The resulting reads were then processed to remove adapters using Porechop v.0.2.4. Finally, the assembler Flye v.2.7.1 (Kolmogorov, Yuan et al. 2019) was used for the assembly of Nanopore MinION reads.

We generated the three primary assemblies using ABySS v.2.2.5 (Simpson, Wong et al. 2009), Supernova v.2.1.1 (Weisenfeld, Kumar et al. 2017), and Flye v.2.7.1 assemblers from Illumina short-reads, 10X Chromium linked reads, and Nanopore long reads, respectively. The completeness and contiguity of all three assemblies were evaluated using BUSCO and Quast, as described previously. The primary assembly generated by Flye showed better assembly metrics and gene completeness compared to those generated by ABySS and Supernova. Consequently, we subjected the Flye assembly to further processing.

Purgehaplotigs (Roach, Schmidt and Borneman 2018) was used to remove redundant and duplicated contigs from the Flye assembly to improve its quality. The haplotigs that were removed during the process were reutilised for scaffolding the assembly with RagTag v.2.1.0 (Alonge, Lebeigle et al. 2021). We utilised filtered long-read data to scaffold and gap-close the assembly with five iterations of Lrscaf v.1.1.11 (Qin, Wu et al. 2019), Rails v.1.5.1 and Cobbler v.0.6.1 (Warren 2016), and Lrgapcloser (Xu, Xu et al. 2019). After each iteration, the resultant assembly was evaluated for completeness and contiguity using Quast. The resulting assembly was further scaffolded with the supernova assembly using RagTag v.2.1.0. We then used the raw 10X linked reads to scaffold the assembly with ArbitR v.0.2 (Hiltunen, Ryberg and Johannesson 2021), Arks v.1.0.4 (Coombe, Zhang et al. 2018) and Links v.1.8.7 (Warren, Yang et al. 2015). We then employed the trimmed paired-end Illumina mRNA sequencing reads (trimming parameters in the section below) to scaffold the assembly with Rascaf (Song, Shankar and Florea 2016). We used Purgehaplotigs to remove haplotigs and then utilised those haplotig reads for two rounds of scaffolding with RagTag.

Blobtools2 (Laetsch and Blaxter 2017) was used to remove the bacterial contaminants. We also excluded small size contigs (<1000 bp) or of a low coverage (<5x coverage) from the assembly and used them after to scaffold the assembly using RagTag. Finally, the assembly was polished with Pilon v.1.24 (Walker, Abeel et al. 2014) using filtered paired-end Illumina shotgun sequencing data to generate the final assembly.

### *De novo* repeat library

We used several *de novo* repeat and homology-based identifiers: LTRharvest (Walker, Abeel et al. 2014), LTRdigest (Steinbiss, Willhoeft et al. 2009), RepeatModeler (Flynn, Hubley et al. 2020), TransposonPSI (Haas, Zeng et al. 2011) (http://transposonpsi.sourceforge.net/) and SINEBase (Flynn, Hubley et al. 2020), to create a custom repeat library for the final *S. obsoletus* genome. Individual libraries from the above steps were concatenated and merged then after. The redundant reads with over 80% of sequence similarity were removed using usearch v.11.0.667 (Edgar 2010). We then used RepeatClassifier to classify the library and mapped reads with unknown categories against UniProtKB/Swiss-Prot database (e-value < 1e-01). Sequences that failed to annotate as repeat sequences were subsequently removed. The final repeat library was used to generate a report of genome repeat content using Repeatmasker v.4.1.2 (Tarailo-Graovac and Chen 2009). The final output was integrated into the MAKER2 (Holt and Yandell 2011) pipeline to mask the genome.

### RNA-sequencing assembly

The mRNA sequencing reads were assessed for quality using FastQC v. 0.10.1 (Andrews 2017) and subjected to further processing to remove Illumina adapters, low-quality bases and short reads. The trimming was performed using Trimmomatic/0.39-Java-1.8.0_144 (Bolger, Lohse and Usadel 2014) with the following settings: TruSeq3-PE-2. fa: 2:30:10 LEADING:3 TRAILING:3 SLIDINGWINDOW:4:15 MINLEN:35. The resulting filtered reads were assembled using Trinity v.2.8.6 (Grabherr, Haas et al. 2011) with the default parameters.

### Genome annotation

We conducted genome annotation using the MAKER2 pipeline (Holt and Yandell 2011) in three iterations, combining evidence-based and ab initio gene model predictions. The first round utilised evidence-based gene model prediction, for which we supplied 269,593 mRNA transcripts *de novo* assembled through the Trinity pipeline (Grabherr, Haas et al. 2011) and 5,281 mRNA and 13,621 protein sequences from Entiminae family obtained from the NCBI database. For the subsequent two rounds, ab initio model prediction was used. Snap (Li, Ma et al. 2007) was trained after each round of MAKER2 v.2.31.9 and used for ab initio gene model prediction on the subsequent runs.

We predicted protein sequences from the MAKER2 pipeline for functional annotation with InterProScan v.5.51-85.0 (Jones, Binns et al. 2014), and retrieved InterPro ID, PFAM domains and Gene Ontology (GO) terms. To assign gene descriptors, we performed top BLASTp hits (Myers, Altschul and Lipman 1990) against the Uniprot database. We assessed the completeness of the predicted proteins and mRNAs using BUSCO.

## Results and discussion

### Genome size estimates

*Sitona obsoletus* genome size was estimated using two different approaches. The first approach involved flow cytometry, which estimated genome size of 1.27 ± 0.01 Gb (mean ± SD). The second involved a k-mer based approach on Illumina shotgun sequencing reads, which estimated the genome size of 0.70 Gb, with unique content of ∼56.4% and a heterozygosity level of 1.68% (Figure 2). The estimated genome size of *S. obsoletus* is comparable to the other forage weevils sequenced, such as *Listronotus bonariensis* with an estimated genome size of 1.1 Gb (Harrop, Le Lec et al. 2020), and *S. discoideus* genome with an estimated size of 0.94 Gb (Katuwal, Bhattarai et al. 2022). Our results showed that estimated genome size of *S. obsoletus* falls within the broad range of coleopteran genomes, which range between 0.25–3.5 Gb.

**Figure 2.**
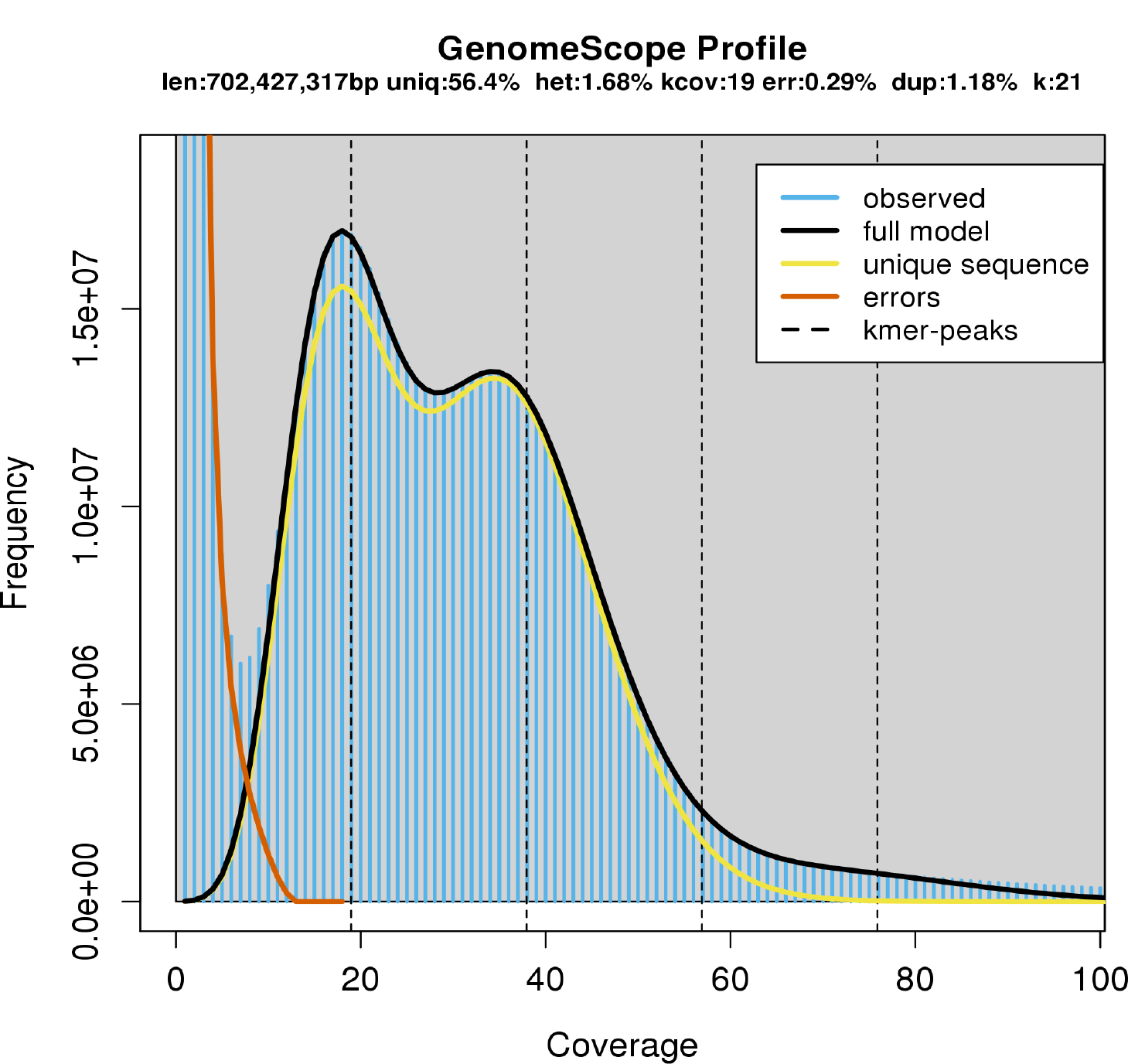
GenomeScope k-mer profile plot for short-read paired-end (PE) libraries illustrating the distribution of 21-mers (k) in 250 bp insert size PE data. K-mer analysis showed the genome size (len) of ∼702 Mb, with 1.68% heterozygosity (het). 56.4% of the genome is unique (uniq) suggesting the rest, 43.6% of the genome as repeats. The mean heterozygous k-mer is at 19-mer (kcov). The error and duplication percentage of the reads are 0.29 and 1.18 respectively.

### Transcriptome assembly

The Trinity pipeline generated an assembly of 223,164,702 bp in total length distributed among 267,052 contigs with an N50 of 4,643 bp. The assembly has GC ratio of 37.75%, and 58,418 contigs were longer than 10,000 bp. An evaluation of the assembly completeness using the insecta_odb10 database with BUSCO (v.5.3.2) showed a BUSCO completeness of 97.9%.

### Genome assembly

Shotgun Illumina short-read sequencing generated 181,173,349 (2x251 bp) raw reads, which underwent the quality control and filtering procedures as previously described. The “clean” reads thus obtained were used for ABySS assembly, resulted in an assembly size of 0.34 Gb spread across 5,077,680 contigs. The N50 and L50 values were 5.52 Kb and 20,190, respectively. The quality of assembly was assessed with Quast, which reported a complete BUSCO of 55.78% and a partial BUSCO of 16.83% from the eukaryotic database. Similarly, sequencing of 10X Genomics Chromium linked reads generated 162.2 million (2x151 bp) paired end reads. These reads were subjected to Supernova assembly, which resulted in an assembly size of 0.26 Gb distributed across 158,370 contigs. The N50 and L50 values were 7.12 Kb and 10,739, respectively (Table 1). The quality of Supernova assembly was evaluated using Quast, which reported a complete BUSCO of 45.54% and a partial BUSCO of 4.62% from the eukaryotic database.

**Table 1.**
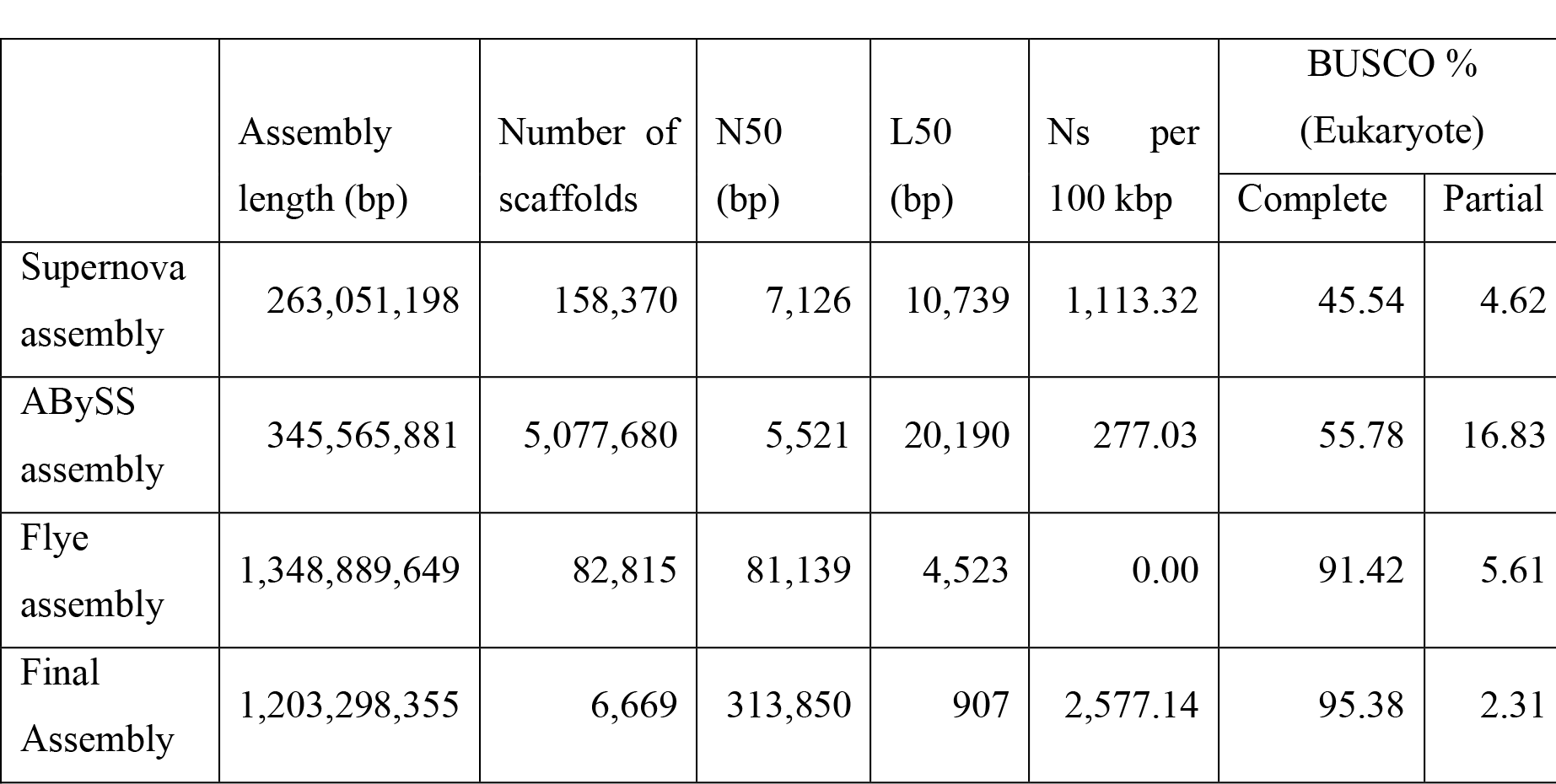
Assembly statistics of *Sitona obsoletus* genome.

In contrast, Nanopore MinION generated over 7 million reads with a total of 9,836,569,248 bases. Quality assessment of the nanopore reads with pycoqc showed a median length of 893 bp, an N50 length of 11,300 bp and a median PHRED score of 12.59 (Supplementary Table 1). The primary assembly from the Flye resulted in an assembly size of 1.34 Gb distributed over 82,815 contigs, with an N50 of 81.14 Kb and L50 of 4,523 (Table 1). The quality of Flye assembly was assessed using Quast, which reported 91.42% of complete BUSCO and 5.61% of partial BUSCO. Notably, the Flye assembly demonstrated significantly better contiguity and assembly statistics compared to the assemblies generated by ABySS and Supernova. Hence, we selected Flye assembly as the primary assembly, while the other assemblies and reads were utilised for scaffolding, gap-closing, and polishing purposes.

The final assembled hybrid genome generated combining short, long and linked read data showed an accumulated length of ∼1.2 Gb. The final assembly consisted of 6,669 scaffolds and an N50 value of 313.85 kb, which represented a significant improvement in assembly contiguity compared to the individual assemblies (Table 1). The L50 value indicated that half of the genome is represented in around 907 scaffolds. Overall, the hybrid assembly showed a 91.94% reduction in the number of scaffolds, a three-fold increase in N50 length, and a 4% increase in BUSCO scores compared to the long-read-only assembly. The BUSCO gene completeness analysis showed that 95.1% of orthologs were present as a complete single copy, with an additional 7.8% were present in duplicated form and 2.5% present in fragmented form (Figure 3). The progress in each assembly statistics after each processing step is given in Supplementary Table 3.

**Figure 3.**
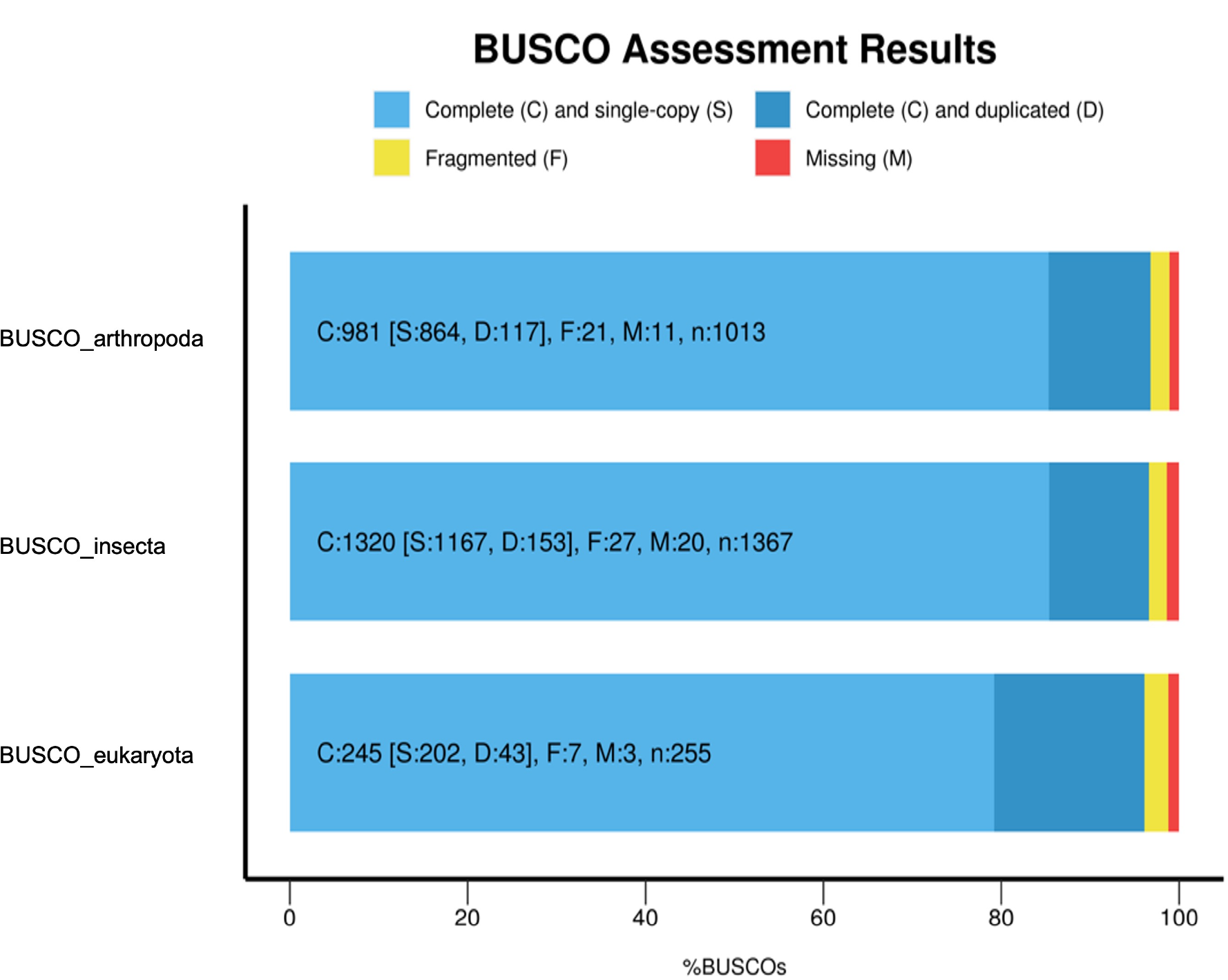
BUSCO v.5 reports for the final hybrid assembly of the *Sitona obsoletus* genome. BUSCO scores in percentage (x-axis) from Arthropoda, Insecta and Eukaryota (odt10) databases (y-axis) are shown in the bar plot. The light blue portion of the bar represents complete and single copy orthologs, dark blue represents complete and duplicated orthologs, yellow represents fragmented orthologs and red represents missing orthologs.

The only other genome sequenced weevil from the genus *Sitona* is *S. discoideus*, which has an N50 of 297.58 kb, with a complete and partial BUSCO score of 96.04% and 1.32% respectively (Katuwal, Bhattarai et al. 2022). Similarly, the only other invasive forage weevil with a reported whole genome from the family Curculionidae is *L. bonariensis*, whose draft genome assembly has an N50 of 122 kb and a complete BUSCO score of 83.9% (Harrop, Le Lec et al. 2020). *Sitona obsoletus*, with an estimated genome size and N50 value of 313.85 kb like *L. bonariensis*, exhibits a higher complete BUSCO score of 95.38%. This suggests that the *S. obsoletus* genome assembled in this study is of particularly high-quality and contiguity in terms of gene completeness. Therefore, it represents a valuable addition as a reference genome for future comparative analysis.

### Genome repeat content

Considering a large distribution of genome sizes in weevils ranging from 162.6 MB to 2,025 MB (Mei, Jing et al. 2022), we also analysed the *S. obsoletus* genome for its repetitive content. Our results showed that 84.26% of the genome was composed of repeats, as determined using Repeat Masker. 64.64% were reported to be the total interspersed repeats, consisting of 22.18% retroelements, 25.04% DNA transposons, 17.19% rolling circles, and 17.41% unclassified elements. Additional repeat categories included small RNA (0.22%), satellites (0.16%), simple repeats (2.01%), and low complexity (0.04%) (see Table 2). The repeat content of other curculionids’ genomes varied, ranging from 81.45% in *S. discoideus* with a genome size of 0.94 Gb (Katuwal, Bhattarai et al. 2022) to 76.36% in *Pachyrhynchus sulphureomaculatus* with a genome size of 2.05 Gb (Van Dam, Cabras et al. 2021), and 70.13% in *L. bonariensis* with a genome size of 1.11 Gb (Harrop, Le Lec et al. 2020). In comparison to these closely related weevils, *S. obsoletus* showed a slightly higher genome repetition, which may account for its slightly larger genome size (1.2 Gb) compared to its close relative *S. discoideus*.

**Table 2.** Repeat content analysis of *Sitona obsoletus* genome.

|  |  |  |  |
| --- | --- | --- | --- |
| Total number of sequences: | 6,669 |  |  |
| Total length: | 1,203,298,355 bp (1,172,319,161 bp excl N/X-runs) |  |  |
| GC level: | 31.84% |  |  |
| Total bases masked: | 1,013,890,500 bp (84.26 %) |  |  |
|  | Number of elements* | Length occupied (bp) | Percentage of sequence (%) |
| Retroelements | 1,065,277 | 266,907,177 | 22.18 |
| SINEs: | 3,164 | 413,456 | 0.03 |
| Penelope | 49,943 | 11,940,546 | 0.99 |
| LINEs: | 736,823 | 151,111,629 | 12.56 |
| CRE/SLACS | 5,246 | 753,639 | 0.06 |
| L2/CR1/Rex | 209,281 | 46,718,465 | 3.88 |
| R1/LOA/Jockey | 17,201 | 5,213,937 | 0.43 |
| R2/R4/NeSL | 3,819 | 1,185,357 | 0.1 |
| RTE/Bov-B | 93,998 | 15,562,933 | 1.29 |
| L1/CIN4 | 1,042 | 158,314 | 0.01 |
| LTR elements: | 325,290 | 115,382,092 | 9.59 |
| BEL/Pao | 25,731 | 12,473,384 | 1.04 |
| Ty1/Copia | 11,361 | 5,564,178 | 0.46 |
| Gypsy/DIRS1 | 147,856 | 66,398,072 | 5.52 |
| Retroviral | 134,177 | 28,720,091 | 2.39 |
| DNA transposons | 1,435,171 | 301,353,269 | 25.04 |
| hobo-Activator | 60,838 | 11,714,999 | 0.97 |
| Tc1-IS630-Pogo | 301,198 | 68,161,954 | 5.66 |
| En-Spm | - | - | 0.00% |
| MuDR-IS905 | - | - | 0.00% |
| PiggyBac | 5,595 | 1288265 | 0.11 |
| Tourist/Harbinger | 16,378 | 3,362,090 | 0.28 |
| Other (Mirage P-element<br>Transib) | 12,038 | 2,843,308 | 0.24 |
| Rolling circles | 1,068,441 | 206,866,386 | 17.19 |
| Unclassified: | 980,690 | 209,528,228 | 17.41 |
| Total interspersed repeats: |  | 777,788,674 | 64.64 |
| Small RNA: | 16,113 | 2,660,405 | 0.22 |
| Satellites: | 10,727 | 1,900,744 | 0.16 |
| Simple repeats: | 184,396 | 24,194,372 | 2.01 |
| Low complexity: | 10,266 | 479,919 | 0.04 |

Note: \* most repeats fragmented by insertions or deletions were counted as one element

### Genome annotation

The assembled and polished genome was annotated using the MAKER2 pipeline, combining both evidence-based and *ab initio* gene models. Following the pipeline, our analysis identified a total of 9,777 genes and 13,521 mRNAs, with a mean gene length of 9,580 bp and a total gene length of 93,668,176 bp, representing 7.78% of *S. obsoletus* assembly (Table 3). Of the predicted annotations, 66% of mRNAs and 65% of proteins were annotated functionally via either one or more of the InterPro, Gene ontology and/or Pfam databases (refer to Supplementary Table 2). The annotated transcriptome had a complete BUSCO score of 64.96%, while the annotated protein achieved a complete BUSCO score of 63.18% when evaluated against the Arthropoda database (refer to Figure 4). Our gene model prediction was had high confidence, with 98% of gene models representing Annotation Edit Distance (AED) scores of 0.5 or less (Supplementary Figure S1).

**Table 3.**
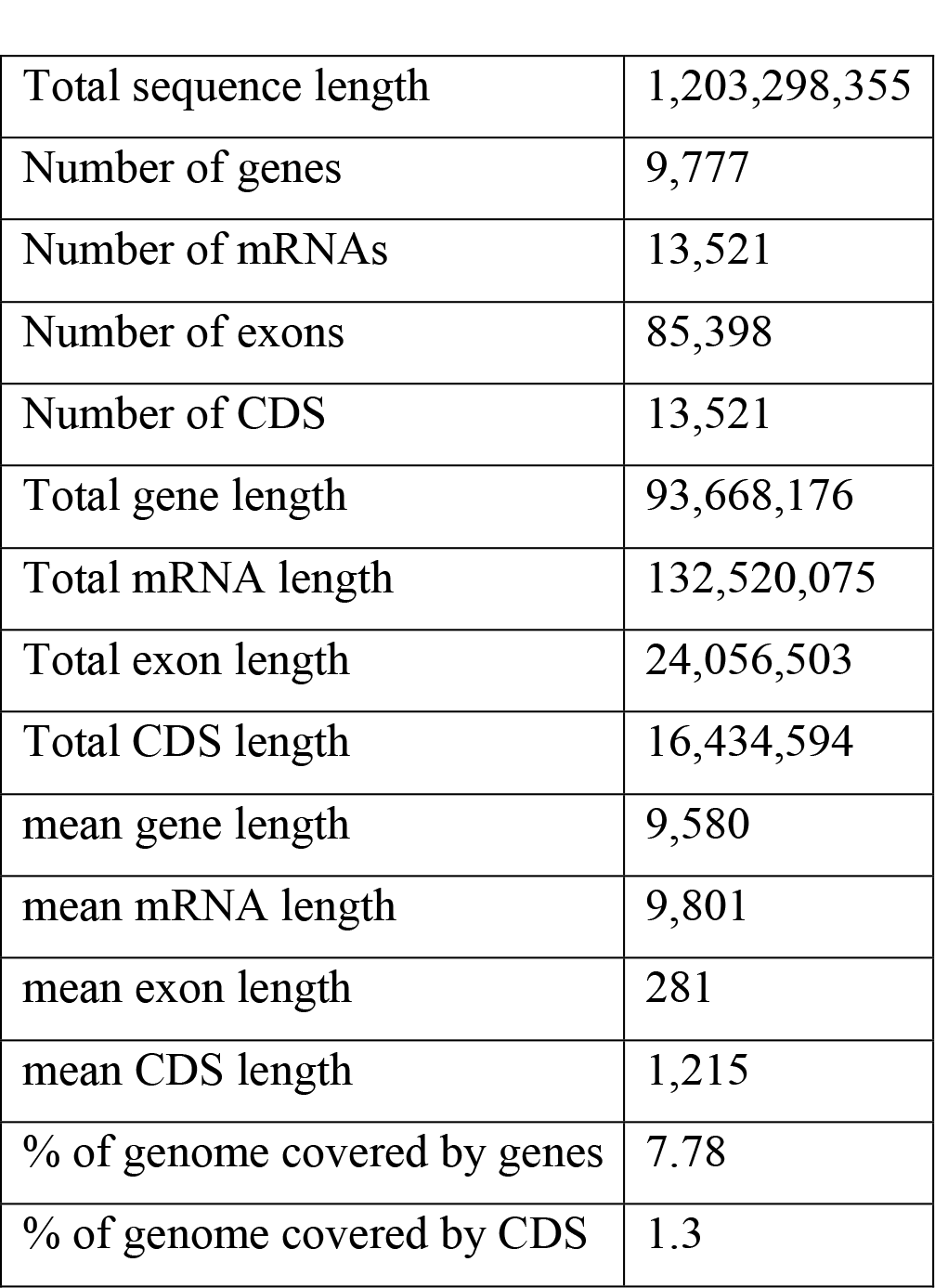
Genome annotation summary for *Sitona obsoletus*.

|  |  |
| --- | --- |
| Total sequence length | 1,203,298,355 |
| Number of genes | 9,777 |
| Number of mRNAs | 13,521 |
| Number of exons | 85,398 |
| Number of CDS | 13,521 |
| Total gene length | 93,668,176 |
| Total mRNA length | 132,520,075 |
| Total exon length | 24,056,503 |
| Total CDS length | 16,434,594 |
| mean gene length | 9,580 |
| mean mRNA length | 9,801 |
| mean exon length | 281 |
| mean CDS length | 1,215 |
| % of genome covered by genes | 7.78 |
| % of genome covered by CDS | 1.3 |

**Figure 4.**
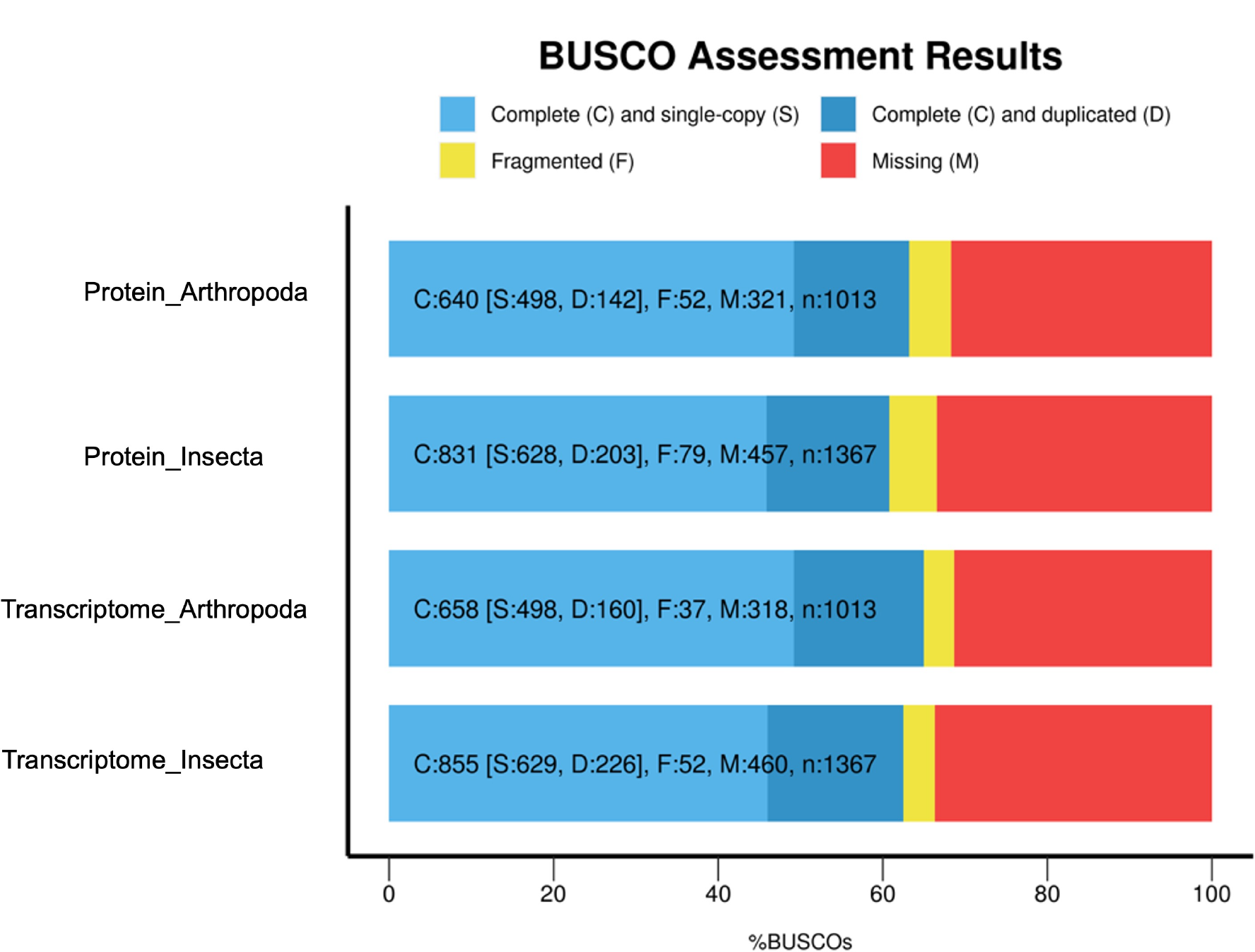
Genome annotation completeness of *Sitona obsoletus* genome through BUSCO database. The plot shows the BUSCO percentage (x-axis) for the annotated Proteins and Transcriptomes using Insecta_odb10 and Arthropda_odb10 database as indicated on the y-axis.

## Conclusion

We utilised a combination of long, linked, and short read technologies to sequence *S. obsoletus* genome. The annotated genome of high-quality and contiguity will be a significant addition to a crucial lineage Coleoptera and will facilitate a range of future studies, including comparative genomics and functional genetics.

## Acknowledgments

We are grateful to Dr. Scott Hardwick (AgResearch, Lincoln), Diane Barton and Colin Ferguson (AgResearch, Invermay) for their assistance in collecting and identifying the weevils. We thank Nicky Richards (AgResearch, Lincoln) for supplying the weevil eggs for laboratory rearing, essential for our RNA-sequencing experiments. Additionally, we acknowledge the support team at New Zealand eScience Infrastructure (NeSI) for granting us access to their high-performance computing platforms, crucial for our data analysis.

