## Supplementary Table S1 for "Whole genome assembly and annotation of the clover root weevil (*Sitona obsoletus*) using a combination of Illumina, 10X Genomics and MinION sequencing"

Supplementary Table S1. Nanopore sequencing output

| Number of Reads | Total Bases | Median Read Length | N50 Length | Median PHRED score |
|---|---|---|---|---|
| 7,285,293 | 23,896,400,000 | 893 | 11,300 | 12.59 |

Supplementary Table S2. Functional annotation statistics from different databases

| Database | No. of terms linked to mRNA | No. of mRNA with term | No. of gene with term |
|---|---|---|---|
| InterPro | 12640 | 8924 | 6354 |
| Gene ontology term | 10827 | 5356 | 3772 |
| Pfam | 12767 | 8924 | 6354 |

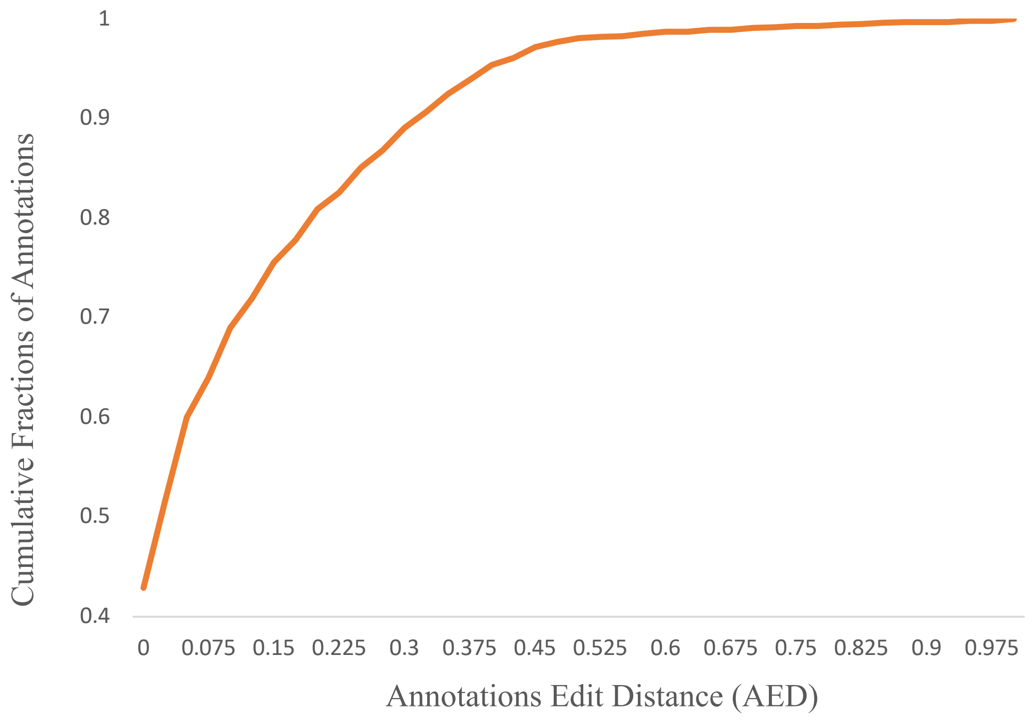


Supplementary Figure S1. Annotation Edit Distance of the predicted genes from three rounds of MAKER2 pipeline.

Supplementary Table S3. Assembly statistics after each round of processing

| | Assembly length | No. of scaffolds | N50 | L50 | Ns per 100 kbp | BUSCO% | |
|---|---|---|---|---|---|---|---|
| | | | | | | Complete | Partial |
| Supernova assembly | 263051198 | 158370 | 7126 | 10739 | 1113.32 | 45.54 | 4.62 |
| Abyss assembly | 345565881 | 5077680 | 5521 | 20190 | 277.03 | 55.78 | 16.83 |
| Flye assembly | 1348889649 | 82815 | 81139 | 4523 | 0.00 | 91.42 | 5.61 |
| Purgehaplotigs | 1141017910 | 51390 | 100826 | 3201 | 0.00 | 90.10 | 6.27 |
| RagTag | 1141395110 | 47618 | 112580 | 2917 | 33.52 | 90.10 | 5.94 |
| Lrscaff | 1506543281 | 26788 | 177442 | 2470 | 8566.22 | 94.72 | 2.64 |
| Rails&Cobbler | 1507291159 | 26741 | 178346 | 2465 | 8276.39 | 94.06 | 2.97 |
| Lrgapcloser | 1507298474 | 26741 | 178249 | 2465 | 3002.56 | 94.06 | 3.3 |
| RagTag | 1507354274 | 26183 | 179559 | 2449 | 3005.40 | 93.73 | 3.63 |
| ArbitR | 1506888250 | 26098 | 181484 | 2418 | 3005.72 | 94.39 | 3.30 |
| Arks & Links | 1507003150 | 24949 | 212032 | 2036 | 3013.16 | 94.39 | 3.30 |
| Rascaf | 1506904542 | 24773 | 216456 | 1980 | 3006.78 | 94.06 | 2.97 |
| Purgehaplotigs | 1245313910 | 18582 | 241252 | 1497 | 3045.76 | 93.73 | 3.3 |
| RagTag | 1245450210 | 17219 | 267645 | 1332 | 3055.98 | 93.40 | 3.3 |
| Blobtools | 1208106584 | 7980 | 272487 | 1280 | 2599.43 | 93.07 | 3.30 |
| RagTag | 1208237684 | 6669 | 315721 | 907 | 2609.90 | 92.74 | 3.63 |
| Pilon | 1203298355 | 6669 | 313850 | 907 | 2577.14 | 95.38 | 2.31 |
